# *Syngap^+/-^* CA1 pyramidal neurons exhibit upregulated translation of long mRNAs associated with LTP

**DOI:** 10.1101/2025.02.19.639128

**Authors:** Aditi Singh, Manuela Rizzi, Sang S. Seo, Emily K. Osterweil

**Affiliations:** Rosamund Stone Zander Translational Neuroscience Center, F. M. Kirby Center, Department of Neurology, Harvard Medical School, Boston Children’s Hospital, Boston, Massachusetts, USA; Simons Initiative for the Developing Brain, University of Edinburgh, UK; Centre for Discovery Brain Sciences, University of Edinburgh, UK

**Keywords:** Syngap, fragile X, LTP, LTD, translation

## Abstract

In the *Syngap^+/-^* model of SYNGAP1-related intellectual disability (SRID), excessive neuronal protein synthesis is linked to deficits in synaptic plasticity. Here, we use Translating Ribosome Affinity Purification and RNA-seq (TRAP-seq) to identify mistranslating mRNAs in *Syngap^+/-^* CA1 pyramidal neurons that exhibit enhanced synaptic stability and impaired long-term potentiation (LTP). We find the translation environment is significantly altered in a manner that is distinct from the *Fmr1^-/y^* model of Fragile X Syndrome (FXS), another monogenic model of autism and intellectual disability (ID). The *Syngap^+/-^* translatome is enriched for regulators of DNA repair, and mimics changes induced with chemical LTP (cLTP) in WT. This includes a striking upregulation in the translation of mRNAs with a longer length (>2kb) coding sequence (CDS). In contrast, long CDS transcripts are downregulated with induction of Gp1 metabotropic glutamate receptor induced long-term depression (mGluR-LTD) in WT, and this profile is mimicked in the *Fmr1^-/y^* model. Together, our results show the *Syngap^+/-^* and *Fmr1^-/y^* models mimic the translation environments of LTP and LTD, respectively, consistent with the dysregulation of these plasticity states in each model. Moreover, we show that translation of >2kb mRNAs is a defining feature of LTP that is oppositely regulated during LTD, revealing a novel mRNA signature of plasticity.

## Background

New protein synthesis in neurons is required to support experience dependent learning and is constitutively altered in multiple mouse models of neurodevelopmental disorders [1–4]. Two notable examples of this are Fragile X Syndrome (FXS) and *SYNGAP1* related intellectual disability (SRID), which arise from mutations in *FMR1* and *SYNGAP1*, respectively [5–9]. Both disorders are commonly identified single-gene causes of autism and ID that co-occur with other behavioral symptoms including hyperactivity, anxiety, and hypersensitivity to sensory stimuli [10, 11]. In the *Fmr1^-/y^* model, excessive protein synthesis in the hippocampus facilitates exaggerated long-term synaptic depression downstream of Gp1 mGluR activation (mGluR-LTD) [5, 6, 12]. Although excessive hippocampal protein synthesis has also been observed in the *Syngap^+/-^*mouse, much less is known about the role of this change in the observed synaptic plasticity phenotypes.

SynGAP is an essential scaffolding protein that modulates the insertion of AMPA-type glutamate receptors at the postsynaptic density (PSD) [13, 14]. Along with its role in PSD complexes, SynGAP also contains a GTP activating (GAP) domain that negatively regulates the activity of small GTPases Ras and Rap1 at synapses [13–15]. The Ras-ERK pathway is a potent regulator of translation, and the excess protein synthesis in *Syngap^+/-^*neurons is corrected with inhibitors of this pathway [8, 9]. A major plasticity deficit observed in *Syngap^+/-^* hippocampus is a significant impairment in LTP induction [14, 16, 17]. Reduction in SynGAP expression in mouse or human cultured neurons results in a persistent increase in AMPARs at the PSD, and increased dendritic spine size indicative of synaptic strength [13, 18]. Extensive in vivo dendritic imaging studies show that SynGAP must be dispersed to allow for insertion of new AMPARs to support LTP [17, 19]. Together, these results suggest that there is a persistent synaptic strengthening in *Syngap^+/-^* that impairs induction of LTP [8, 14]. In addition to this described role in LTP, a study investigating mGluR-LTD in the *Syngap^+/-^* mouse also revealed an exaggeration similar to the *Fmr1^-/y^* model [9].

Here, we sought to understand the role of altered protein synthesis in the hippocampal plasticity phenotypes seen in the *Syngap^+/-^* mouse. To do this, we performed Translating Ribosome Affinity Purification and RNA-seq (TRAP-seq) to profile translating mRNAs in the CA1 pyramidal neurons of *Syngap^+/-^* hippocampus [20]. We find dysregulation of a number of transcripts, including a surprising upregulation in those encoding DNA regulatory proteins and chromatin modifiers. We also find that there is little overlap between the differentially expressed transcripts in *Syngap^+/-^* and *Fmr1^-/y^* mutant models. Interestingly, changes seen in the *Syngap^+/-^*TRAP are similar to those induced in WT slices with chemical stimulation of LTP (cLTP), assessed by comparing to previously-published CamK2a-ribotag data from cLTP-stimulated hippocampal slices [21]. In contrast, there is little overlap between *Syngap^+/-^* TRAP and mGluR-LTD TRAP populations. Indeed, a comparison of TRAP-seq datasets from WT stimulated for cLTP or mGluR-LTD reveals a striking divergence in these opposing plasticity states. Gene ontology (GO) and Gene set enrichment analysis (GSEA) reveal an increased translation of axonal and cell adhesion effectors during cLTP, and a decrease in these factors during mGluR-LTD. In contrast, increased translation of ribosomal and mitochondrial proteins is induced with mGluR-LTD, and these are decreased during cLTP. Further investigation reveals an opposite regulation of translating mRNA populations based on transcript coding sequence (CDS) length. Shorter-length (<1kb) transcripts encoding metabolic regulators including ribosomal and mitochondrial proteins are reduced, and longer-length (>2kb) transcripts encoding synaptic and cell adhesion proteins are increased, during cLTP. The opposite length-dependent shift is seen with induction of mGluR-LTD. The same opposite relationship is seen in *Syngap^+/-^* and *Fmr1^-/y^* models. Together, our results show the translating mRNA population in *Syngap^+/-^* CA1 mimics that induced by cLTP in WT, including the upregulation of long mRNAs, which may contribute to the persistent synaptic strengthening in this model.

## Results

### *Syngap^+/-^* and *Fmr1^-/y^* CA1 translatomes are largely dissimilar

The identity of the overly-synthesized protein population in *Syngap^+/-^* neurons is not known, and we therefore performed TRAP-seq on *Syngap^+/-^* and WT littermates expressing EGFP-L10a in CA1 pyramidal neurons as in previous work (**Fig. 1A**) [22, 23]. Our results show that 145 transcripts are differentially expressed in the *Syngap^+/-^*CA1- TRAP fraction (*P*adj < 0.1) (**Fig. 1B, Additional file 1**). Of these mRNAs, 69 are upregulated and 76 are downregulated. In contrast to the ribosome-bound TRAP fraction, a comparison of WT and *Syngap^+/-^* total RNA fractions reveals only a small number of differentially expressed transcripts, with 7 significantly upregulated and 12 significantly downregulated (*P*adj < 0.1) (**Fig. 1B, Additional file 2**). The most significantly changed transcripts in the TRAP fraction do not exhibit a similar change in the total RNA fraction, suggesting the effects are not solely due to transcript abundance (**Fig. S1**).

**Figure 1.**
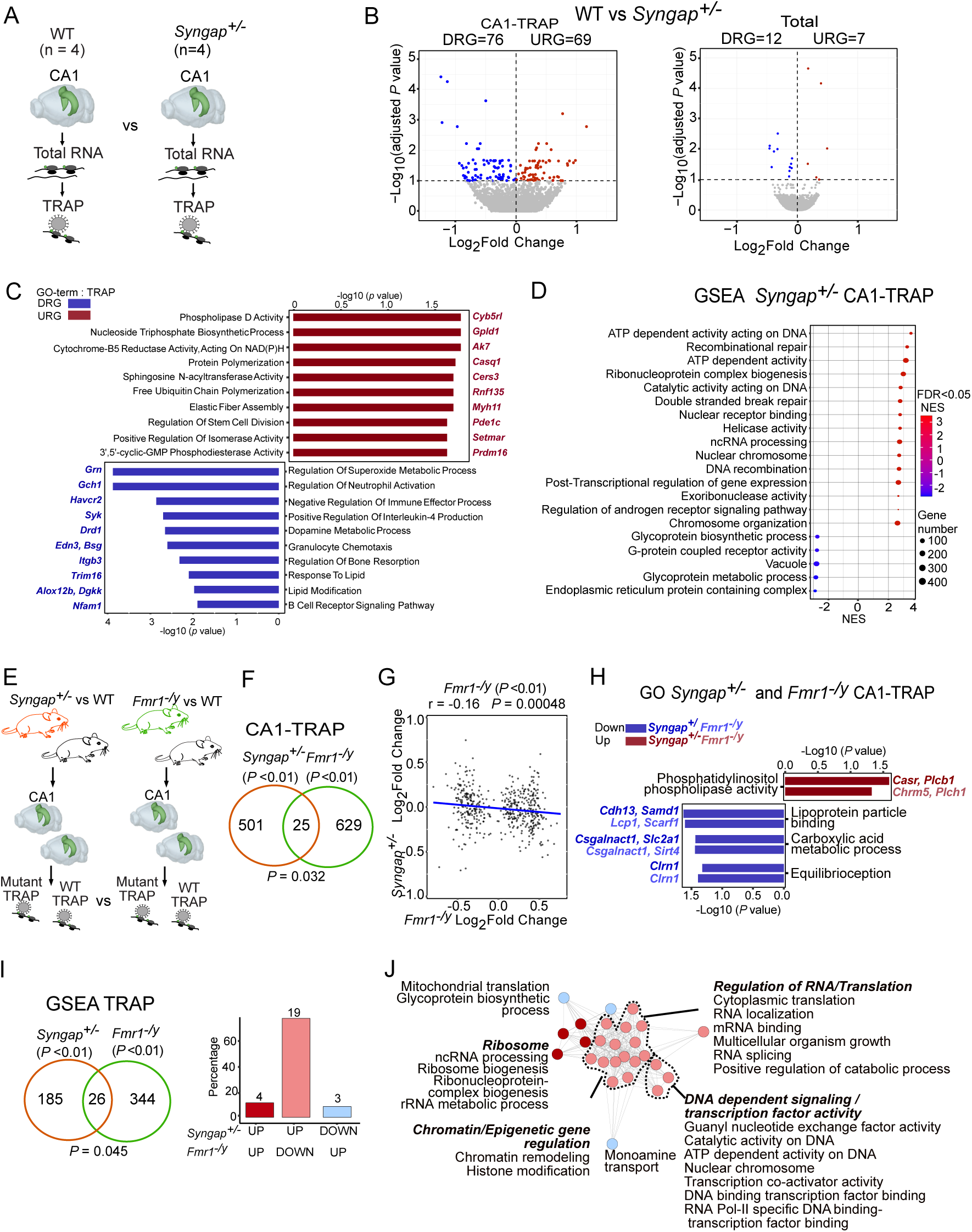
The translating mRNA population in *Syngap^+/-^* CA1 neurons is enriched for DNA repair proteins, and is distinct from the population in *Fmr1^-/y^* CA1 neurons. (A) Schematic for TRAP-seq and Total RNA-seq analysis of *Syngap^+/-^*vs WT (N = 4 littermate pairs) from hippocampal CA1 neurons. (B) Volcano plots for differential analysis of TRAP-seq data on the left and Total RNA-seq on the right show substantial changes in translatome than the transcriptome of *Syngap^+/-^*. Significant transcripts (adjusted *P* value < 0.1) being under-translated or under-expressed are denoted in blue and over-translated or over-expressed are denoted in red. (C) Gene ontology (GO) analysis of transcripts upregulated and downregulated in translatome between *Syngap^+/-^* vs WT (D) Gene set enrichment analysis (GSEA) of the *Syngap^+/-^* (FDR<0.05) shows a significant downregulation of glycoprotein metabolism, endoplasmic reticulum and vacuole related activities while upregulation of processes related to DNA repair, recombination, including ATP-dependent activity acting on DNA and transcriptional regulation. (E) Schematic for TRAP-seq dataset comparison of *Syngap^+/-^* vs WT to the TRAP-seq of *Fmr1^-/y^*vs WT from hippocampal CA1 neurons. (F) Quantification of transcripts shows 526 significant transcripts (*P* value < 0.01) are differentially translating in *Syngap^+/-^* and only 25 of those overlap with the significant translatome in *Fmr1^-/y^*(\**P* = 0.032). (G) Transcripts significantly changed in *Fmr1^-/y^*translatome are negatively correlated with *Syngap^+/-^* translatome changes (r = −0.16, \**P* =0.00048). (H) Gene ontology (GO) analysis of transcripts significantly (*P* value < 0.01) altered in *Syngap^+/^* and *Fmr1^-/y^* shows only few functional processes are regulated similarly in both mutants (*P* value < 0.05). (I) To determine whether the gene sets altered in *Syngap^+/-^*are similar to those altered in the *Fmr1^-/y^* translating population, significantly changed gene sets (adjusted *P* value < 0.01) were compared to those significantly changed in the *Fmr1^-/y^* population (adjusted *P* value < 0.01). This reveals a modest overlap of 25 gene sets (\**P* = 0.045) but 19 of these (85%) are regulated in opposite direction. (J) The gene sets inversely modulated between *Syngap^+/-^*and *Fmr1^-/y^* regulate the chromatin remodeling, RNA localization and metabolism, transcription factor and DNA dependent activities among others, which are upregulated in *Syngap^+/-^* while downregulated in *Fmr1^-/y^*.

To understand how the differentially expressed transcripts alter molecular and cellular process in *Syngap^+/-^*, we performed a GO analysis on the most significantly up- and downregulated populations (*P*adj < 0.1). This revealed an upregulation in categories related to cytochrome-B5 reductase activity (acting on NAD(P)H), nucleoside triphosphate biosynthetic process, and phospholipase D activity which suggests an increase in cellular functions associated with energy metabolism, nucleotide synthesis, and neuronal membrane remodeling. Additionally, the enrichment of positive regulation of isomerase activity, free ubiquitin chain polymerization, sphingosine N-acyltransferase activity, and protein polymerization points to increased protein modification and elongation, potentially contributing to the observed phenotype of altered protein synthesis (**Fig. 1C, Additional file 3**). The downregulated population is enriched for immune-related signaling pathways, dopamine metabolic process, and regulation of lipid modification suggesting a reduction in neurotransmitter metabolism. To complement these findings and gain a comprehensive understanding of functional enrichment - capturing gene sets with subtle, coordinated changes rather than focusing solely on the most significantly altered genes - we next performed Gene Set Enrichment Analysis (GSEA) on the CA1-TRAP population. This showed upregulation of processes related to DNA modification, including recombinational repair and ATP-dependent activity (led by *Mcm2 and Mcm4* core enrichment), as well as RNA regulatory terms suggesting increased genomic stability and transcriptional regulation demands in *Syngap*^+/-^ neurons (**Fig. 1D**, **Additional file 4**). In contrast, the most significantly downregulated gene sets are related to endoplasmic reticulum, glycoprotein metabolism and vacuoles. In neurons, these pathways are critical for protein folding, trafficking, and synaptic signaling. Together, these changes highlight a potential trade-off in *Syngap*^+/-^ neurons, with increased investment in DNA repair and stability at the expense of glycoprotein metabolism and receptor activity. This imbalance may impact cellular resilience, signaling efficiency, and ultimately synaptic function, potentially contributing to the impaired cognitive and behavioral phenotypes.

Both *Syngap^+/-^*and *Fmr1^-/y^*mice are models of autism and ID, however they express different plasticity phenotypes in hippocampal CA1. In the *Fmr1^-/y^* mouse, a basal elevation of protein synthesis occludes further translation downstream of mGluRs, resulting in an exaggeration of mGluR-LTD that no longer requires protein synthesis [5, 6, 24–26]. Although an increase in mGluR-LTD is seen in the *Syngap^+/-^*hippocampus, there is also a robust deficit in LTP that arises from saturation of AMPA receptors at the postsynaptic density that prevents further potentiation [19]. To investigate whether similar differences were seen in the translating mRNA populations, we compared *Syngap^+/-^* CA1-TRAP to *Fmr1^-/y^* mice bred to the same CA1-TRAP line (**Fig. 1E**) [22, 23]. A comparison between *Syngap^+/-^*versus *Fmr1^-/y^*CA1-TRAP populations (*P*adj<0.1) is not significant (*P*=0.052) with only one common transcript (**Fig. S2).** Since we are comparing two distinct TRAP datasets, we used a slightly relaxed threshold (*P*<0.01), which reveals a small but significant 4.75% overlap of differentially expressed transcripts (\**P* = 0.032), however only 15 mRNAs are dysregulated in the same direction (**Fig. 1F, Additional file 5**). Additionally, acorrelation between significantly altered transcripts in the *Fmr1^-/y^*CA1-TRAP with the expression of these genes in *Syngap^+/-^* CA1-TRAP reveals a small but significant negative correlation (\**P* = 0.00048) (**Fig. 1G**). Furthermore, even with a relaxed threshold (*P* value < 0.05), the GO analysis of significantly altered transcripts in the *Fmr1^-/y^* CA1-TRAP shows minimal overlap with the GO terms identified in *Syngap^+/-^* (also at *P* value < 0.05). The only overlapping terms are related to membrane remodeling, metabolism, and equilibrioception (**Fig. 1H, Additional file 6**). This indicates a lack of similarity between the two mutant models.

To assess whether there was any similarity in the gene sets altered in *Syngap^+/-^* and *Fmr1^-/y^* mutants, we compared GSEA results from the TRAP fractions from each mutant. Our results show that there is a 12% significant overlap (\**P* = 0.045) (**Fig. 1I, Additional file 7**). However, the vast majority of overlapping terms (84.6%) are changed in the opposite direction. The terms shared in both mutants relate to chromatin, transcription, DNA activity and RNA splicing, which are upregulated in the *Syngap^+/-^*CA1-TRAP and downregulated in the *Fmr1^-/y^*CA1-TRAP (**Fig. 1J**). Together, these results suggest that shared changes in the *Syngap^+/-^* and *Fmr1^-/y^* models are mostly opposing.

### *Syngap^+/-^*, but not *Fmr1^-/y^* CA1-TRAP shows changes consistent with cLTP

In our previous study, we performed TRAP-seq on CA1 pyramidal neurons in acute hippocampal slices 30 min after application of a 5 minute pulse of 50 µM S-DHPG, which induces robust mGluR-LTD [23]. We compared these changes to those basally altered in *Fmr1^-/y^* CA1-TRAP and found a similarity that suggests a saturation of LTD-related protein synthesis. Given the opposite profiles seen between *Syngap^+/-^*and *Fmr1^-/y^*CA1-TRAP populations, and the saturation of LTP in *Syngap^+/-^* CA1, we wondered whether the mistranslating population in *Syngap^+/-^* would be similar to that of LTP induction in WT. To investigate this, we analyzed a dataset generated by Chen et al., who performed ribotag pulldown and RNA-seq on Camk2a-positive CA1 and CA3 neurons in acute hippocampal slices 30 minutes post-stimulation with 50 µM forskolin to induce robust cLTP (**Fig. 2A**) [21]. The RNA-seq reads for LTP datasets were re-mapped to the current Ensemble mouse genome and processing approach similar to *Syngap^+/-^* and *Fmr1^-/y^* datasets (See Methods). Then, control versus stimulated populations were compared using DESeq2 with parameters identical to all datasets (*P*adj < 0.1).

**Figure 2.**
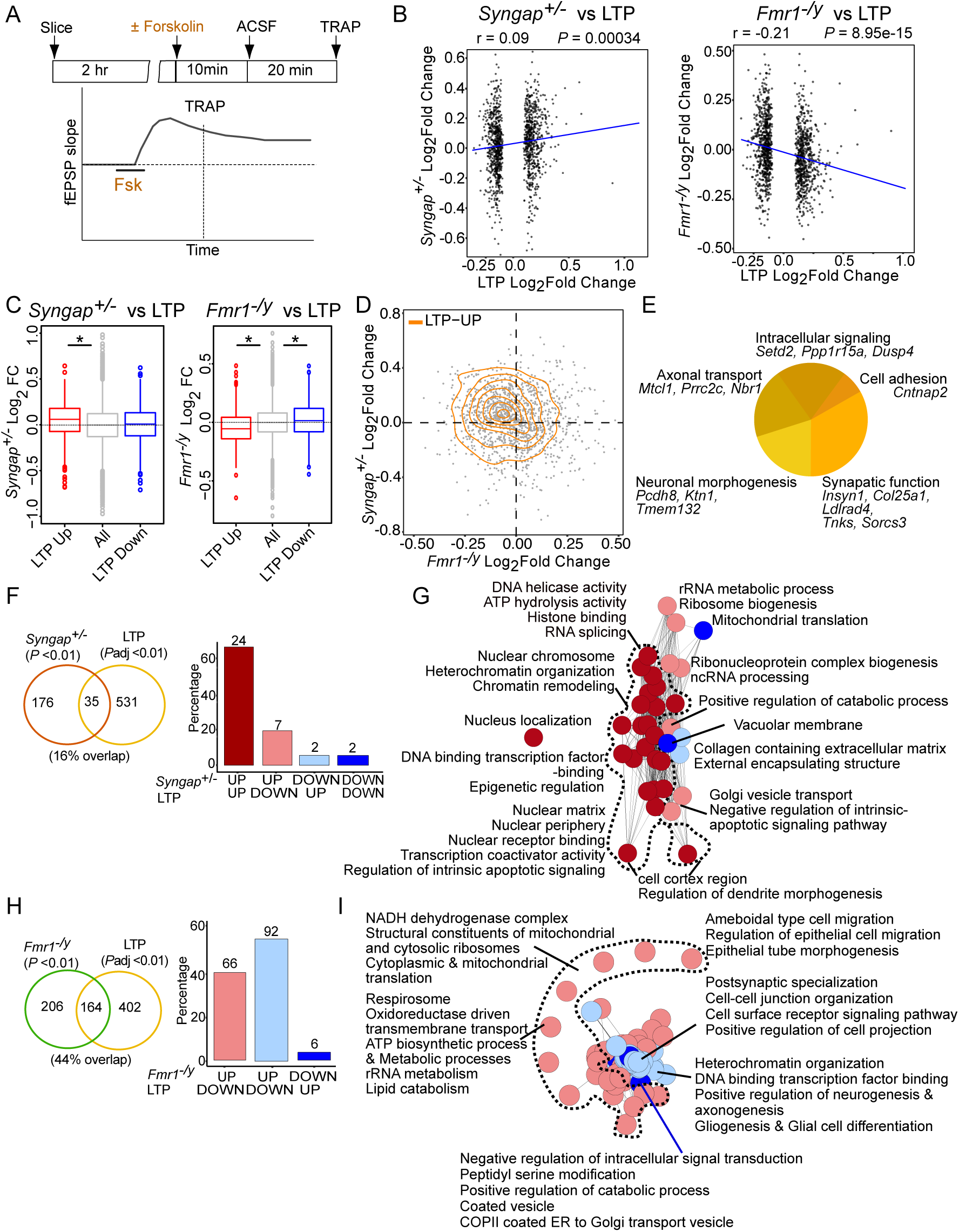
cLTP-specific translation changes in WT match basal changes in *Syngap^+/-^* CA1 neurons, but diverge from changes in *Fmr1^-/y^* CA1 neurons. **(A)** Schematic of the TRAP strategy from wild type (WT) hippocampal slices stimulated with 50 µM forskolin to induce robust cLTP chemical LTP (Chen et al) followed by ribotag pulldown and RNA-seq on Camk2a-positive CA1 and CA3 neurons. **(B)** LTP-specific significant transcripts (adjusted *P* value < 0.1) show small but significantly positive correlation with *Syngap^+/-^* translatome (r = −0.09, \**P* =00034) while notably negative correlation with *Fmr1^-/y^* translatome changes (r = −0.21, \**P* =8.95e-15). **(C)** Analysis of the LTP-specific significant transcripts in the *Syngap^+/-^* translatome shows significant increase in LTP- upregulated transcripts but no change in LTP- downregulated transcript (Kruskal-Wallis test \**P* = 2.15e-12, Post hoc two-sided Wilcoxon rank-sum test up \**P* = 2.96e-13, down *P* = 0.055), while LTP-specific significant transcripts in the *Fmr1^-/y^* translatome show significant opposing change both groups-LTP- upregulated and LTP- downregulated transcripts (Kruskal-Wallis test \**P* < 2.2e- 16, Post hoc two-sided Wilcoxon rank-sum test up \**P* < 2.2e-16, down \**P* = 0.00078). **(D)** Joint distribution analysis of LTP-specific transcripts between *Syngap^+/-^* and *Fmr1^-/y^* translatomes in a 2D density plot shows the positive distribution pattern of LTP upregulated transcripts in *Syngap^+/-^*. **(E)** Analysis of the significantly upregulated LTP-specific transcript population that are also upregulated in *Syngap^+/-^* translatome fraction identifies transcripts, which are involved in synaptic functions, intracellular signaling, neuronal morphogenesis, axonal transport and cell adhesion. **(F)** To determine whether the gene sets altered in *Syngap^+/-^*are also altered in the cLTP translating population, significantly changed *Syngap^+/-^*gene sets (*P* value < 0.01) were compared to those significantly changed in the cLTP population (adjusted *P* value < 0.01). This unveils an overlap of 35 gene sets (*P* = 0.054), nonetheless majority of these (74.28%) are similarly upregulated in both. **(G)** The gene sets that are alike and upregulated in both *Syngap^+/-^*and cLTP are involved in dendrite morphogenesis, chromosome organization, transcription and DNA dependent regulatory activities among others. **(H)** Comparison of the gene sets altered in cLTP (adjusted *P* value < 0.01) with the ones altered in *Fmr1^-/y^* (*P* value < 0.01) shows a greater overlap of 164 (44.3%) terms (\**P* = 1.24e-57) however 56% of these terms are changed in an opposite direction. **(I)** Shared gene sets between cLTP and *Fmr1^-/y^* regulate important processes such as mitochondrial function, different metabolic processes and translation which is upregulated in *Fmr1^-/y^* while downregulated with cLTP. The gene sets involved in axonogenesis, synaptic adhesion, postsynaptic specialization and heterochromatin organization are upregulated with LTP but downregulated in *Fmr1^-/y^*.

To rule out changes that occur with general plasticity stimulation, we removed transcripts that were also significantly changed in our LTD dataset, resulting in a “LTP-specific” population. Our results show that there is a significant positive correlation between LTP-induced transcripts and those changed in *Syngap^+/-^* CA1-TRAP (\**P* = 0.00034) (**Fig. 2B, Additional file 8**). In contrast, a similar comparison to the *Fmr1^-/y^* revealed that transcripts upregulated with LTP were significantly negatively correlated (\**P* = 8.95e-15). To ask whether significantly up- and down-regulated populations might be distinctly changed in the *Syngap^+/-^*or *Fmr1^-/y^*populations, we performed a separate analysis to individually compare these groups (**Fig. 2C**). Our results show a significant increase in LTP-upregulated transcripts in the *Syngap^+/-^*(\**P* = 2.96e-13), but no significant downregulation in the LTP-downregulated population (*P* = 0.055). In *Fmr1^-/y^*, there is a significant opposing change observed in both groups (up \**P* < 2.2e-16, down \**P* = 0.00078) (**Fig. 2C**). Furthermore, this trend was consistent across all transcripts that were significantly altered in LTP, not limited to specific populations (**Fig. S3**). These results suggest that the differential translating mRNAs in *Syngap^+/-^*CA1 show similar changes during cLTP. The most significantly upregulated transcripts in LTP that are also upregulated in *Syngap^+/-^* include regulators of synaptic function and plasticity (*Insyn1, Col25a1, Ldlrad4, Tnks, Sorcs3*), neuronal morphogenesis (*Pcdh8, Ktn1, Tmem132*), axonal transport (*Mtcl1, Prrc2c, Nbr1*), intracellular signaling (*Setd2, Ppp1r15a, Dusp4*) and cell adhesion (*Cntnap2*) (**Fig. 2D-E**).

Next, we used GSEA to compare gene sets significantly changed in *Syngap^+/-^* CA1-TRAP with those changed during cLTP. Our results show that 17% of gene sets changed in the *Syngap^+/-^*population are also changed with cLTP in WT (*P* = 0.054) (**Fig. 2F, Additional file 9**). Importantly, most of the overlapping terms are shifted in the same direction (74.28%). The majority of similarly upregulated categories include those involved in chromosome organization, transcription and DNA regulation (**Fig. 2G**). In stark contrast, a comparison between LTP and *Fmr1^-/y^*populations shows a 44.3% overlap (\**P* = 1.24e-57), however a remarkable 56% of these terms are changed in an opposite direction (**Fig. 2H**). The overlapping categories upregulated in *Fmr1^-/y^* and downregulated with LTP include those involved in mitochondrial function and ribosomes (**Fig. 2I**). Categories upregulated with LTP and downregulated in *Fmr1^-/y^* include those involved in synaptic adhesion and heterochromatin organization. These results indicate that changes to the translating mRNA population of hippocampal pyramidal neurons induced by cLTP are similar to basal changes in *Syngap^+/-^* hippocampus and opposite to basal changes in *Fmr1^-/y^*hippocampus.

### *Fmr1^-/y^*, but not *Syngap^+/-^* CA1 pyramidal neurons show translation changes similar to those induced with mGluR-LTD

Despite the basal synaptic strengthening and LTP occlusion seen in *Syngap^+/-^* hippocampus and cortex, a previous study also revealed an exaggeration of hippocampal mGluR- LTD in in this model [9, 14, 17]. We therefore compared the expression profile seen in CA1-TRAP isolated from WT slices 30 min after stimulation of mGluR-LTD (published in [23], reprocessed with identical parameters- see methods) to the profile seen in *Syngap^+/-^*CA1-TRAP (**Fig. 3A**). Transcripts overlapping with the LTP dataset were removed, resulting in a “LTD-specific” population. We find no correlation between transcripts changed with LTD and those changed in *Syngap^+/-^* neurons (*P* = 0.53) (**Fig. 3B, Additional file 10**). This contrasts with the small but significant positive correlation between LTD and *Fmr1^-/y^* populations (\**P* = 1.37e-06). A comparison of significantly up- and downregulated populations shows a small but significant downregulation of transcripts upregulated with LTD in the *Syngap^+/-^* population (\**P* = 2.741e-05), and no change in the LTD-downregulated population (**Fig. 3C**). In contrast, the LTD Up- and down-regulated populations are changed in a similar direction in the *Fmr1^-/y^*pool (Up *\*P* = 7.049e-09, Down *P* = 0.05777) (**Fig. 3C-D**). The trends from LTD-specific transcripts are replicated even for total significantly altered transcripts *Syngap^+/-^* and *Fmr1^-/y^* (**Fig. S3**). A functional breakdown of mRNAs significantly upregulated with LTD that are also upregulated in *Fmr1^-/y^* highlights those involved in ribosome function (*Rpl9, Rps25, Rpl41*), cellular transport (*Tma7, Tmsb46, Chmp1a*), Cytokine production (*Alox5, Litaf, Trim56*), mitochondria (*Pole4, Atp5j*), apoptosis (*Ier3ip1, Dynll1*), oxidative stress (*Mt1*), and transcription (*Zic1*) (**Fig. 3E**).

**Figure 3.**
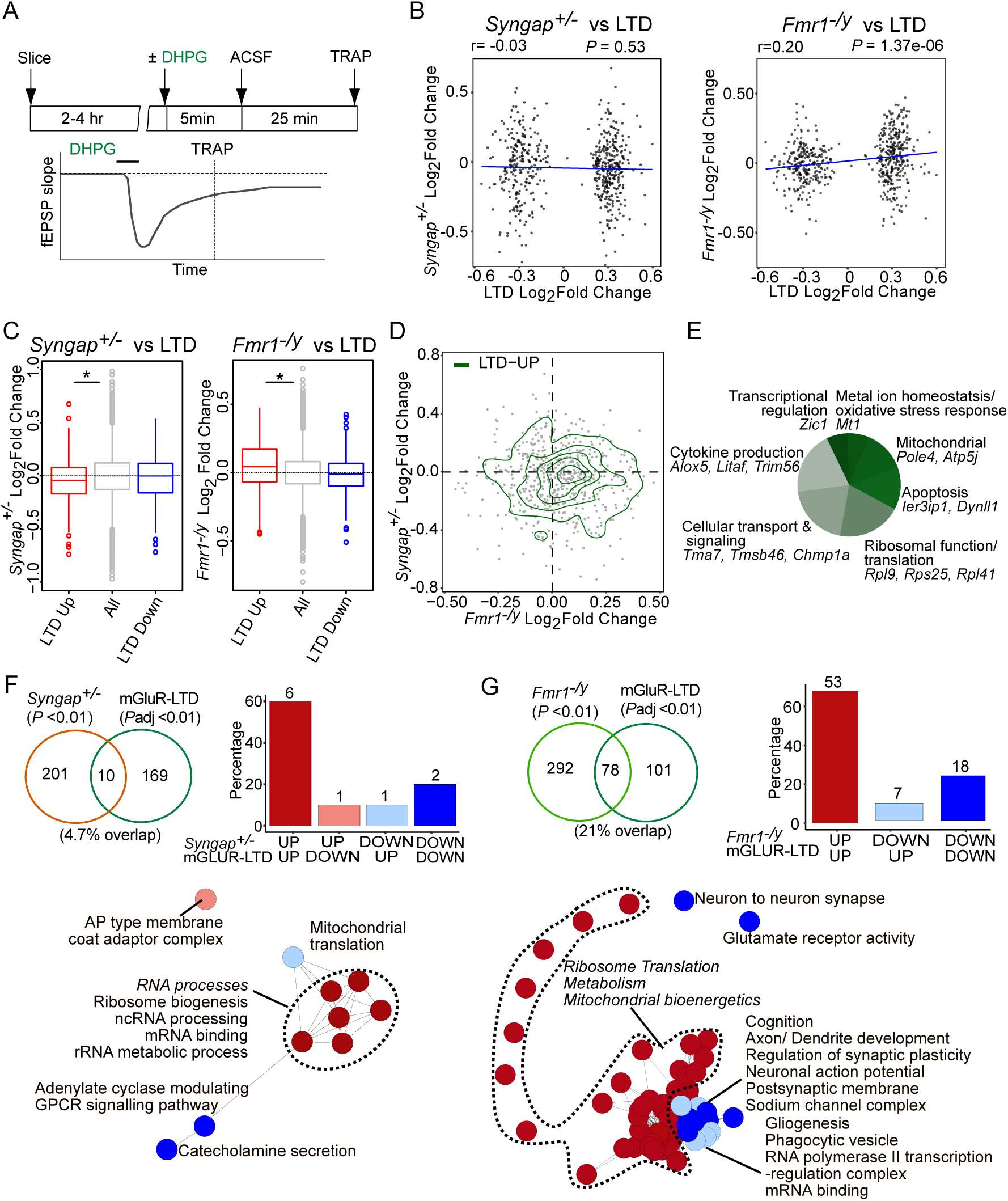
mGluR-LTD specific translation changes in CA1 neurons match basal changes *Fmr1^-/y^* but not *Syngap^+/-^* hippocampus. **(A)** Schematic of the TRAP strategy from wild type (WT) hippocampal slices stimulated with a 5 minute pulse of 50 µM S-DHPG that induces robust mGluR- LTD (Seo et al) followed by TRAP-seq on CA1 pyramidal neurons. **(B)** LTD-specific significant transcripts (adjusted P value < 0.1) show no correlation with *Syngap^+/-^* translatome (r = −0.03, *P* =0.533) while remarkably significant positive correlation with *Fmr1^-/y^* translatome changes (r =0.20, \**P* =1.375e-06). **(C)** Analysis of the LTD-specific significant transcripts in the *Syngap^+/-^* translatome shows significant decrease in LTD- upregulated transcripts (Kruskal-Wallis test \**P* = 0.0001, Post hoc two-sided Wilcoxon rank-sum test up \**P* = 2.74e-05, down *P* = 0.07). In contrast, LTD-specific significant transcripts in the *Fmr1^-/y^* translatome show notably significant increase in LTD- upregulated transcripts (Kruskal-Wallis test \**P* =2.66e-08, Post hoc two-sided Wilcoxon rank-sum test up \**P* =7.04e-09, down *P* = 0.057). **(D)** Combined distribution analysis of LTD-specific transcripts between *Syngap^+/-^* and *Fmr1^-/y^* translatomes in a 2D density plot shows the positive distribution pattern of LTD upregulated transcripts with *Fmr1^-/y^* translation changes. **(E)** Analysis of the significantly upregulated LTD-specific transcript population that are also upregulated in *Fmr1^-/y^* translatome fraction identifies transcripts which are involved in apoptosis, ribosomal as well as mitochondrial functions, transcription regulation and cellular transport. **(F)** Comparison of the gene sets significantly altered in *Syngap^+/-^* (*P* value < 0.01) with the ones altered in LTD (adjusted *P* value < 0.01) reveals non-significant overlap of merely 4.7% (*P* = 0.3927527). **(G)** To determine whether the gene sets altered in mGluR-LTD translating population match *Fmr1^-/y^* translatome, significantly changed *Fmr1^-/y^*gene sets (*P* value < 0.01) were compared to those significantly changed in the LTD population (adjusted *P* value < 0.01). This unveils a greater overlap of 21% with 78 terms (\**P* = 1.83e-46) and 91% of these terms are changed in similar direction. The gene sets that are up-regulated alike in LTD and *Fmr1^-/y^*are involved in ribosome and mitochondrial function, while similarly downregulated sets are related to neuronal activity and synaptic membrane functions.

Next, to investigate any similarities in functional groups we compared gene set enrichment in LTD and *Syngap^+/-^* populations. This revealed a small 4.7% overlap between *Syngap^+/-^* and LTD populations, with only 8 changed in a similar direction (*P* = 0.39) (**Fig. 3F, Additional file 11**). In contrast, a comparison between LTD and *Fmr1^-/y^*reveals a 21% overlap with 78 gene sets shifted in the same direction (\**P* = 1.83e-46) (**Fig. 3G**). The similarly upregulated gene sets are those relating to ribosome and mitochondrial function, and similarly downregulated sets are related to neuronal and synaptic function. These results suggest that the translation profile induced with LTD is similar to the basal population in *Fmr1^-/y^*but not *Syngap^+/-^*CA1.

### The mRNA populations translated during LTP and LTD are largely divergent

Our *Syngap^+/-^*comparisons led us to the realization that there are significant differences between the translating mRNA populations induced with LTP versus LTD in hippocampal neurons. Although there have been many differential expression studies examining the differences in gene expression between LTP and LTD, we are not aware of a direct comparison between translation profiles in CA1 pyramidal neurons. We therefore compared these populations (**Fig. 4A**). Interestingly, a comparison of significantly changed transcripts revealed most were stimulation-specific (1280 LTP, 593 LTD), however a significant overlap of 105 transcripts was also observed (*\*P* = 1.14e-12) (**Fig. 4B, Additional file 12**). Many of the shared transcripts upregulated in both stimulations are immediate early genes (IEGs) (**Fig. 4C**) that mark neuronal activation (i.e., *Fos, Npas4, Junb*, etc.) [27, 28]. To assess the functional relevance of transcripts significantly upregulated with cLTP, we performed GO analyses (**Fig. 4D, Additional file 13**). This shows an enrichment in axon development, dendrite development, and synapse structure. In contrast, the population significantly upregulated with induction of mGluR-LTD is enriched for cytoplasmic translation, respiration, and metabolism (**Fig. 4E**). These results indicate the transcripts translated during cLTP and those translated during mGluR-LTD encode for very different functional classes. The shared population upregulated by both stimulations is enriched for transcription factors and regulators of membrane excitability including IEGs, while the shared downregulated population is enriched for calcium regulators and GTPases (**Fig. S4**). This is consistent with a general cellular response to stimulation.

**Figure 4.**
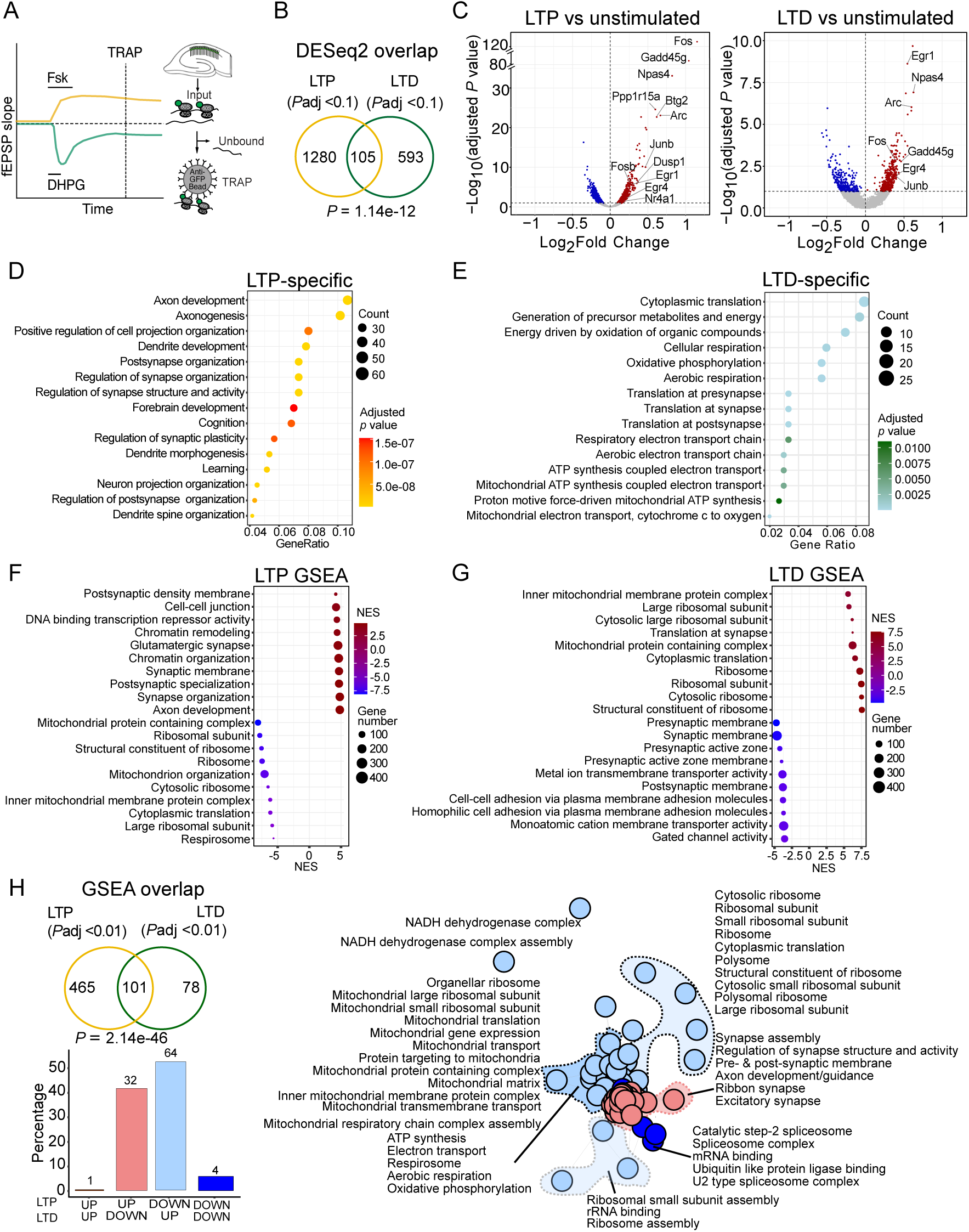
LTP and LTD induce distinct translatome shifts including opposite changes in synaptic transcripts. **(A)** Schematic of the TRAP strategy from wild type (WT) mouse hippocampal slices induced for mGluR-LTD (Sang et al) and chemical LTP (Chen et al). **(B)** LTP and LTD comparison shows distinctly translated transcripts in both phenomena. Differential analysis was performed to identify synaptic plasticity related transcripts i.e. LTP vs unstimulated control and LTD vs unstimulated control. LTP and LTD significant (adjusted *P* value < 0.1) transcripts were compared to find a common overlap and specific population of transcripts. **(C)** Volcano plot of ribosome bound translating population of transcripts in LTP and LTD. Significant transcripts (adjusted *P* value < 0.1) being under-translated are denoted in blue and over-translated are denoted in red. Both LTP and LTD stimulation exhibit upregulated translation of immediate early genes such as *Arc*, *Fos, Npas4* and *Egr1* indicating neuronal activation. LTP specific translation of *Ppp1r15a*, *Btg2* and *Nr4a* among other transcripts is also remarkable. **(D-E)** Gene ontology analysis of LTP and LTD specific transcripts shows their unique functions. LTP specific transcripts predominantly regulate axonogenesis, cell projection and dendrite development processes among others (D) while LTD specific transcripts primarily regulate cytoplasmic translation, pre- & post-synaptic translation, metabolism, and energy precursors synthesis (E). **(F)** GSEA analysis of the LTP vs unstimulated control TRAP-seq dataset identified over translation of gene sets related to synapse organization, cell-cell junction, and chromatin remodeling in LTP (adjusted *P* value < 0.1). The downregulated gene sets in LTP are involved in mitochondrial and ribosomal functions as well as cytoplasmic translation. **(G)** GSEA analysis of the LTD vs unstimulated control TRAP-seq dataset identified over translation of gene sets related to ribosome, translation and mitochondrial terms in LTD (adjusted *P* value < 0.1), while the downregulated gene sets are involved in pre and post synaptic membrane functions as well as cell-cell adhesion. **(H)** To determine uniquely altered gene sets altered, a comparison of significant (adjusted *P* value < 0.1) gene sets identified by GSEA in both LTP and LTD TRAP-seq datasets was performed. This reveals a common pool of 101 gene sets (\**P* = 2.14e-46) and their remarkably opposite regulation between LTP vs LTD. The gene sets involved in synapse assembly, axon development and regulation of synapse structure are up- in LTP but downregulated in LTD. The gene sets related to cytoplasmic and mitochondrial ribosome, ATP synthesis and electron transport among other significant terms are down regulated in LTP but upregulated in LTD.

To assess whether the gene sets shifted with cLTP and mGluR-LTD were similarly divergent, we performed GSEA on each population. Our analysis of LTP-specific changes revealed an upregulation of synaptic terms and transcription/chromatin regulators, and a downregulation of ribosome- and mitochondria-related terms (**Fig. 4F, Additional file 14**). Conversely, LTD-specific changes include an upregulation of ribosome- and mitochondria-related terms and a downregulation of synaptic terms (**Fig. 4G**). Interestingly, a comparison of gene sets shifted with each stimulation showed a significant overlap (101 shared, 465 LTP, 78 LTD; *\*P* =2.146771e-46), but a striking divergence in the direction of change within shared categories (**Fig. 4H**). Specifically, LTP is defined by a significant upregulation in synaptic stability transcripts and a downregulation in ribosomal and mitochondrial transcripts. In contrast, LTD is defined as an upregulation in ribosomal/mitochondrial transcripts and a downregulation in synaptic stability gene sets. Together, these results provide compelling evidence that the mRNAs translated to support LTP and LTD are divergent, and a subset is oppositely regulated.

### Long transcripts encoding synaptic structural components are bi-directionally translated with LTP and LTD in hippocampal pyramidal neurons

In previous work, we noted a negative correlation between differential expression and transcript length in the translating population of *Fmr1^-/y^* CA1 pyramidal neurons [23]. This can be seen as a significant increase in shorter mRNAs within the population (i.e., <1kb) and a significant downregulation of the longer transcripts (i.e., >2kb). Interestingly, there is a functional segregation of effectors encoded by genes of differing length within the neuronal genome, that is also seen in the transcriptome [29–31]. In particular, shorter genes encode ribosomal proteins, mitochondrial proteins, nucleosome proteins, and regulators of metabolic function. In contrast, longer genes encode proteins involved in cell adhesion, ion channels, and cytoskeleton proteins. This effect can be seen in total transcript length, but also in the length of the CDS that does not include untranslated regions (UTRs). We hypothesize the length-dependent shift in the neuronal translatome of *Fmr1^-/y^* neurons is contributing to the constitutive underproduction of synaptic stability proteins [23].

To examine whether a similar length-dependent translation shift is present in *Syngap^+/-^*CA1 neurons, we compared differential expression to CDS length in the significantly changed population (*P* < 0.01). Our results show a significant positive correlation between expression and transcript CDS length in the *Syngap^+/-^* TRAP-seq population (*\*P* = 2.2e-16) (**Fig. 5A**). Further analysis shows a significant increase in the length of the upregulated population that does not extend to total transcript length, 3’ UTR length or 5’UTR (**Fig. S5, Additional file 15**). To assess whether this effect can be seen in the entire population, we compared the differential expression of all transcripts binned by CDS length (<1kb, 1-2kb, 2-4kb, >4kb) as compared to the average population. Consistent with our correlation analysis, we find a positive length shift in the *Syngap^+/-^* translatome that can be seen as a reduction in <1kb (*\*P* = 1.669e-14) and an increase in 2-4kb (*\*P* = 2.943e-15) and >4kb (*\*P* < 2.2e-16). As shown previously, this length shift is negative in the *Fmr1^-/y^* TRAP-seq population (**Fig. 5B**).

**Figure 5.**
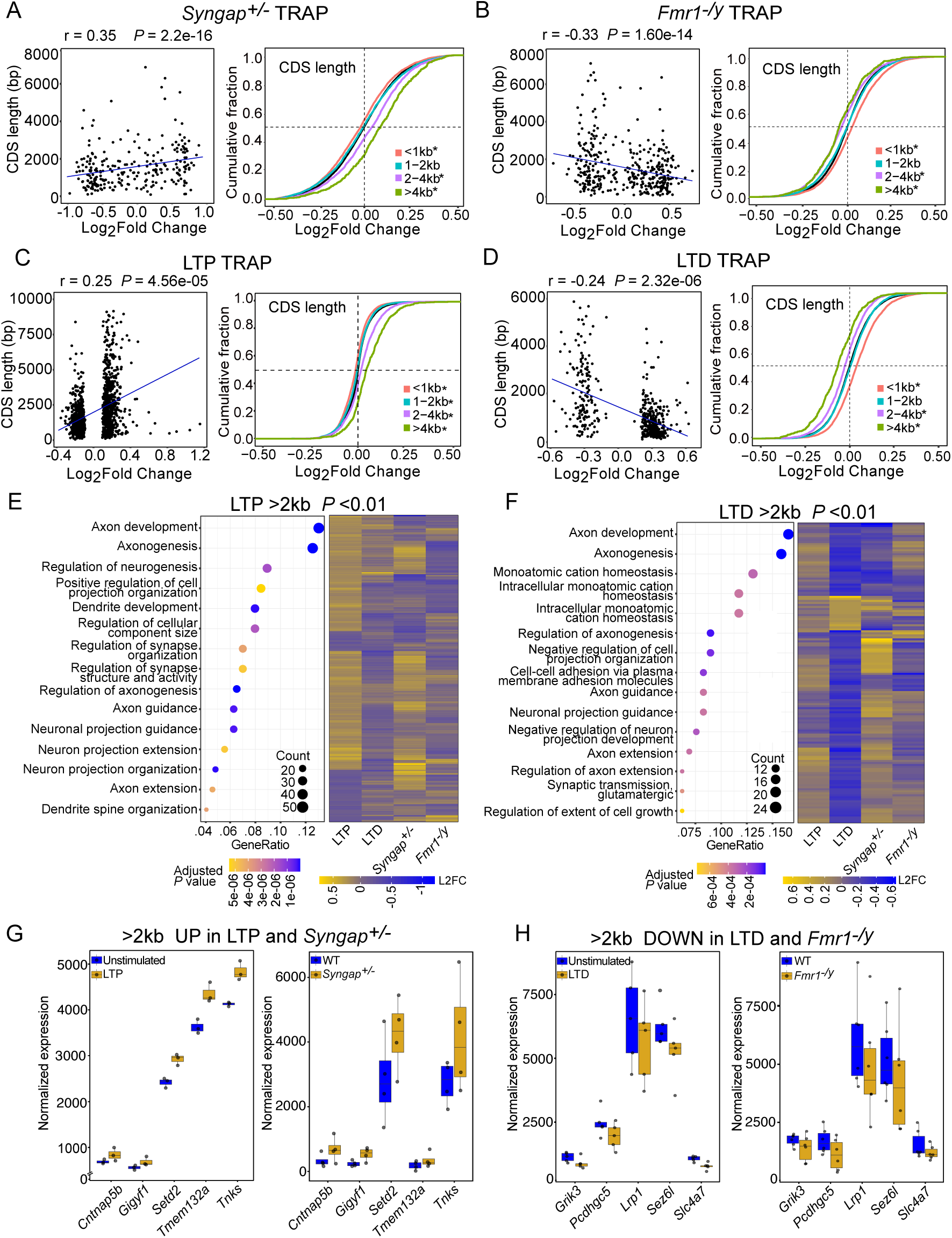
Translation of long (>2kb) transcripts is bi-directionally altered by stimulation of LTP versus LTD, and this is mimicked in *Syngap^+/-^* and *Fmr1^-/y^*mutant CA1 neurons. **(A)** Transcripts significantly changed in *Syngap^+/-^* translatome (P value < 0.01) show significant positive correlation with longer CDS length (left, r = 0.35, \**P* =2.2e-16). A binned analysis on CDS lengths of altered translatome shows that *Syngap^+/-^* TRAP fraction exhibits upregulated translation of longer transcripts (two-sample z test; >4kb vs all: z = 10.716, \**P* <2.2e-16, 2-4kb vs all: z = 7.8933, \**P* =2.94e-15, 1-2kb vs all: z=-0.92171, *P*=0.35, <1kb vs all: z=-7.6739, \**P*=1.66e-14). **(B)** Transcripts significantly changed in *Fmr1^-/y^* translatome (*P* value < 0.01) show significant negative correlation with longer CDS length (left, r = −0.33, \**P* =1.60e-14). A binned analysis on CDS lengths of altered translatome shows that *Fmr1^-/y^*TRAP fraction exhibits decreased translation of longer transcripts (two-sample z test; >4kb vs all: z = −7.5046, \**P* =6.16e-14, 2-4kb vs all: z = −10.079, \**P* <2.2e-16, 1-2kb vs all: z=-1.5061, *P*=0.132, <1kb vs all: z=12.301, \**P*<2.2e-16). **(C)** Analysis of cLTP translatome (*P* value < 0.01) shows significant positive correlation with longer CDS length (left, r = 0.25, \**P* =4.56e-05). A binned analysis on CDS lengths of altered translatome shows that cLTP causes increased translation of longer transcripts (two-sample z test; >4kb vs all: z = 13.987, \**P* <2.2e-16, 2-4kb vs all: z =10.401, \**P* <2.2e-16, 1-2kb vs all: z=-4.9587, \**P*=7.09e-07, <1kb vs all: z=-13.1, \**P* <2.2e-16). **(D)** Analysis of mGluR-LTD translatome (*P* value < 0.01) shows significant negative correlation with longer CDS length (left, r = −0.24, \**P* =2.32e-06). A binned analysis on CDS lengths of altered translatome in LTD exhibits decreased translation of longer transcripts (two-sample z test; >4kb vs all: z = −14.638, \**P* <2.2e-16, 2-4kb vs all: z = −13.203, \**P* <2.2e-16, 1-2kb vs all: z=1.2064, *P*=0.22, <1kb vs all: z=21.175, \**P*<2.2e-16). **(E)** Gene ontology analysis of long transcripts (CDS length >2kb) in LTP shows their functions in axon development, cell projection and synapse structure organization (left). These transcripts are upregulated (log2foldchanges-L2FC >0) and show largely similar alteration with *Syngap^+/-^* while opposite patterns of alteration (L2FC <0) in LTD as well as *Fmr1^-/y^* (right). **(F)** Gene ontology analysis of long transcripts (CDS length >2kb) in LTD shows they are related to axon development, cell projection and synapse organization (left) similar to LTP but translation of these transcripts is decreased in LTD and *Fmr1^-/y^* opposite to LTP and *Syngap^+/-^* (right). **(G-H)** Bidirectionally altered long transcripts in LTP and *Syngap^+/-^* (G) vs LTD and *Fmr1^-/y^* (H).

As the *Syngap^+/-^*population exhibits changes consistent with LTP, we next investigated whether a length-dependent shift exists in the WT population stimulated for cLTP. Our results reveal a striking positive correlation between transcript CDS length and differential expression in the WT ribotag population stimulated for cLTP (*\*P* = 4.56e-05) (**Fig. 5C**). As reported in Chen et al., a significant increase in 3’UTR length is also seen in the upregulated population, along with an increase in total transcript length and 5’UTR length (**Fig. S5, Additional file 16**). The CDS length shift is also observed in a binned analysis across the translatome (<1kb *\*P* < 2.2e-16, 1-2kb *\*P* = 7.096e-07, 2-4kb *\*P* < 2.2e-16, >4kb *\*P* < 2.2e-16). The same analysis of the LTD population reveals a negative correlation (as previously published) (**Fig. 5D**). Together, these results show a positive length-dependent translation shift in *Syngap^+/-^* CA1 neurons that matches that seen with induction of cLTP in WT. This stands in stark contrast to the negative length-shift seen in *Fmr1^-/y^* CA1 neurons and in those induced for mGluR-LTD.

Given the similar upregulation of longer (>2kb) mRNAs in the *Syngap^+/-^* population and in the LTP-induced population, we wondered whether we could observe a profile consistent with persistent synaptic strength in this population. To evaluate this, we performed a GO analysis of the significantly changed transcripts >2Kb in the LTP dataset, the majority of which are upregulated (76%, 321/421) (**Fig. 5E, Additional file 16**). This revealed a clear enrichment in axon and dendrite development as well as synaptic structure, consistent with synaptic strengthening. A heatmap comparing these 421 transcripts shows both the opposing expression in LTD and *Fmr1^-/y^*populations, and a similar expression in the *Syngap^+/-^* population. Next, we investigated the >2kb transcripts significantly changed with mGluR-LTD, the majority of which are downregulated (86.8%, 138/159) (**Fig. 5F**). GO analysis shows that this population is similarly enriched for terms related to axon development and synaptic structure, consistent with a profile for synaptic weakening. A heatmap comparing these 159 transcripts shows both the opposing expression in LTP and *Syngap^+/-^*populations, and a similar expression in the *Fmr1^-/y^* population.

Collectively, our results suggest a model whereby increased translation of long (>2Kb) mRNAs in hippocampal pyramidal neurons, many of which are constitutively upregulated in *Syngap^+/-^*CA1, supports synaptic strengthening. In contrast, reduced translation of long mRNAs, which are constitutively under-translating in *Fmr1^-/y^*, supports synaptic weakening. We therefore identified the most significantly upregulated >2kb transcripts in both *Syngap^+/-^* and LTP populations: *Gigyf1, Cntnap5b, Tnks, Setd2,* and *Tmem132a* (**Fig. 5G**). All of these genes encode regulators of synaptic signaling and stability, and three are candidates that have identified variants linked to ASD (see **Additional file 17**). Transcripts most significantly downregulated in both *Fmr1^-/y^* and mGluR- LTD populations include *Pcdhgc5, Sez6l, Lrp1, Grik3*, and *Slc4a7* (**Fig. 5H**). Four of these genes have variants linked to ASD, and all are involved in synaptic function and excitability (**Additional file 17**). Together, these up- and downregulated targets, majority of which are linked to autism and ID, underscore distinct molecular mechanisms through which *Syngap*^+/-^ and *Fmr1*^-/y^ mouse models show impaired neurodevelopment, synaptic function, and cognitive outcomes.

## Discussion

This study sought to identify differentially translating mRNAs in CA1 pyramidal neurons of the *Syngap^+/-^*hippocampus that might participate in synaptic phenotypes. We find that there is a significant increase in DNA regulators and a correlation with changes induced with cLTP in WT. This is opposite to the translating population seen in *Fmr1^-/y^* CA1, where there is an increase in ribosomal proteins and changes that mimic induction of mGluR-LTD in WT. Interestingly, we also find that cLTP induces a translation profile that is strikingly different from that induced with mGluR-LTD. This includes an increase in the translation of longer mRNAs >2kb, a profile that matches basal changes in the *Syngap^+/-^*population. In contrast, long mRNAs are decreased with induction of LTD in WT and basally downregulated in the *Fmr1^-/y^* population. The >2kb transcripts significantly upregulated upon induction of LTP and downregulated upon induction of LTD encode regulators of axon/dendrite stability and synaptic adhesion. Overlapping the population changed with plasticity with the populations changed in *Syngap^+/-^*or *Fmr1^-/y^*CA1-TRAP identifies a common list of candidates for the occlusion of LTP or LTD in these models.

There are limitations to this study. First, although we use TRAP as a proxy for the translating mRNA population, it is important to note that this population represents a combined measurement of RNA abundance and ribosome association. This means that changes we observe cannot be attributed to translation alone, and may be due to either an increase in ribosome engagement or a change in the availability of mRNA. However, we note the same length-dependent change in *Fmr1^-/y^* neurons is seen in ribosome profiling experiments from fly and mouse models where transcript abundance is not a co-factor [32, 33]. Another limitation is that the paradigms used to stimulate LTP and LTD in hippocampal slices are chemical agonists of PKA or Gp1 mGluRs, which likely have many effects beyond those relevant to synaptic plasticity [21, 23]. Although we cannot know which changes are directly responsible for the change in synaptic strength, we note that there is a similar increase in IEGs that mark neuronal activity in both stimulations (**Fig. 4C**). This suggests that the opposing changes we observe are not likely due to a change in overall cellular activity.

Although there have been several studies of translation differences in mouse models of FXS, relatively few have been performed in *Syngap^+/-^* models. Here, we identify a unique translation signature that could support persistent changes in neuronal function and plasticity. Indeed, our TRAP-seq results reveal an increase nucleotide enrichment of genesets including minichromosome maintenance (MCM) subunits *Mcm2* and *Mcm4* within the *Syngap^+/-^* population (**Fig. 1D**). The MCM complex is a collection of DNA helicases that are essential for unwinding DNA during replication [34, 35]. The association of MCM with DNA maintains genome integrity during cell-cycle progression, and it is essential for controlling the speed of replication [34, 36]. The increase of MCM subunits in the *Syngap^+/-^* TRAP is curious, and suggests a potential mechanism for the precocious neurogenesis phenotype that has been observed in organoids cultured from SYNGAP1 haploinsufficiency patients [37]. It is also possible the MCM upregulation is reflecting an increase in DNA repair, which is associated with neurons undergoing synaptic strengthening [38, 39]. Indeed, overlapping the most significantly changed gene sets in cLTP and *Syngap^+/-^* CA1- TRAP populations reveals similar upregulation in DNA and chromatin regulatory processes (**Fig. 2F-G, Additional file 9**).

Exaggerated, protein synthesis-independent mGluR-LTD has been seen in the *Syngap^+/-^*hippocampus [9], which contrasts with the molecular similarities we observe between the *Syngap*^+/-^ translation profile and that of induced cLTP. Our examination of the *Syngap^+/-^* CA1 translatome does not reveal any obvious similarities to the mGluR-LTD stimulated translatome and instead shows a profile more aligned with LTP. Nonetheless, we cannot rule out the potential contribution of excessive protein synthesis to the *Syngap^+/-^*phenotype. An alternative explanation for this discrepancy may lie in altered AMPA receptor (AMPAR) dynamics in *Syngap*^+/-^ synapses, where the persistent synaptic strengthening and increase in AMPARs at the PSD in *Syngap^+/-^*CA1 neurons results in a greater loss of AMPARs from the postsynaptic membrane upon mGluR stimulation resulting in exaggerated LTD. Consequently, even though the molecular profile of *Syngap^+/-^* CA1 neurons aligns with cLTP, the structural and receptor composition changes in *Syngap^+/-^* synapses may facilitate an exaggerated mGluR-LTD response. In this context, elevated AMPAR availability could prime *Syngap*^+/-^ neurons for heightened synaptic weakening upon mGluR-LTD induction, resulting in an LTD response that is amplified despite the cLTP-like translation profile. Further experiments are necessary to tease apart this mechanism.

Along with the changes in DNA regulators, the transcripts convergently upregulated in *Syngap^+/-^*and cLTP-stimulated WT include regulators of synaptic function and cell adhesion (**Fig. 2E**). Further analyses revealed that both the *Syngap^+/-^* CA1 neurons and those stimulated for cLTP exhibit a significant increase in long (>2kb) mRNAs in the translating fraction (**Fig. 5A,C**). This is opposite to a reduction in >2kb transcripts we observe in *Fmr1^-/y^* CA1 pyramidal neurons and those stimulated for mGluR-LTD in WT (**Fig. 5B,D**). Importantly, this >2kb population is enriched for regulators of synaptic function and cell adhesion (**Fig. 5E-F**), which is consistent with the inherent association of gene length and cellular function that has been described in the neuronal genome [23, 29]. We previously hypothesized that the reduced translation of long transcripts in *Fmr1^-/y^*neurons is proximal to increased ribosome abundance, which unequally impacts translation of mRNAs as a function of length [23]. Others have suggested that the reduced translation of long mRNAs is due to impaired stability of long mRNAs in the absence of FMRP [40–42]. While we do not know the cause of the increased translation of long mRNAs in the *Syngap^+/-^*CA1-TRAP, there are significant changes in RNA binding proteins that may be involved in RNA stability including *Rbm38*, *Rbm47*, *RbmS3*, *Tent2*, and *Pabpc4*. It is also worth noting that DNA repair mechanisms are particularly relevant for longer genes, and the upregulation of MCM may selectively stabilize the transcription of these genes in *Syngap^+/-^* neurons [43, 44].

Our comparison of LTP and LTD datasets reveals a striking opposite regulation of transcripts with a long CDS length in the translating ribosome fraction of CA1 pyramidal neurons. It is possible that this phenomenon represents a specific regulation of transcripts involved in synaptic function. It is tempting to speculate that mechanisms to bias translation toward or away from long transcripts could be a type of gain control that allows specific plasticity states to be supported. Although our experiments do not distinguish between local/dendritic changes versus somatic changes, it is not unreasonable to suggest that such changes could be present in the local environment. Future experiments dissecting the local translating population during LTP or LTD stimulation would be particularly enlightening for answering this question.

## Methods

### Animals

*Syngap^+/-^* mice were originally generated by Komiyama et al (2002) and were a generous gift from Peter Kind and Seth Grant. These mice were bred using heterozygous crosses and maintained on the C57Black6JOla line (Harlan). CA1-TRAP mice (created by http://gensat.org/ and obtained from Jackson Labs with permission from Nathanial Heintz) were bred on the JAX C57BL/6J background. All experiments were carried out using male littermate mice aged P25-32, and studied with the experimenter blind to genotype. *Syngap^+/-^*and WT littermates were bred from a F1 cross of *Syngap^+/-^* females and CA1-TRAP homozygous males. Mice were group-housed (six maximum) in conventional non-environmentally enriched cages with unrestricted food and water access and a 12 h light–dark cycle. Room temperature was maintained at 21 ± 2 °C with ambient humidity. Animal husbandry was carried out by University of Edinburgh technical staff. All procedures were performed in accordance with ARRIVE guidelines and the UK Animal Welfare Act, and were approved by the Animal Welfare and Ethical Review Body at the University of Edinburgh.

### TRAP

TRAP was performed on *Syngap^+/-^*CA1-TRAP littermates as described previously in Thomson et al. 2017. Briefly, male littermates (P25-32) were decapitated and hippocampi rapidly dissected in ice cold PBS. Hippocampi were homogenized in ice-cold lysis buffer (20 mM HEPES, 5 mM MgCl2, 150 mM KCl, 0.5 mM DTT, 100 mg/ml cycloheximide, RNase inhibitors and protease inhibitors) using dounce homogenizers, and samples centrifuged at 1,000 x g for 10 min to remove large debris. Supernatants were then extracted with 1% NP-40 and 1% DHPC on ice, and centrifuged at 20,000 x g for 20 min. A 50 mL sample of supernatant was removed for use as Input, and the rest incubated with streptavidin/protein L-coated Dynabeads (Life Technologies) bound to anti-GFP antibodies (HtzGFP-19F7 and HtzGFP-19C8, Memorial Sloan Kettering Centre) overnight at 4^°^C with gentle mixing. Anti-GFP beads were washed with high salt buffer (20 mM HEPES, 5 mM MgCl2, 350 mM KCl, 1% NP-40, 0.5 mM DTT and 100 mg/ml cycloheximide) and RNA was eluted from all samples using Absolutely RNA Nanoprep kit (Agilent) according to the manufacturer’s instructions. RNA yield was quantified using RiboGreen (Life Technologies) and RNA quality was determined by Bioanalyzer analysis.

### RNA-Seq library preparation and analysis

RNA with RIN > 7 was prepared for RNA-seq using the RNaseq Ovation V2 kit (Nugen), according to manufacturer’s instructions. Samples were sent to Oxford Genomics Centre for sequencing using Illumina HiSeq 2500 or HiSeq 4000. Adapters were removed using cutadapt 2.6 [45] with Python 3.6.3 and fastqc module (version 0.11.9) was used to analyse the quality of reads. Sequencing reads (50 or 75 bp, paired end) from, *Syngap^+/-^ Fmr1^-/y^*, cLTP and mGLUR-LTD datasets were mapped to Mus musculus primary assembly (Ensembl release v109) of Mouse Genome GRCm39 using STAR (Spliced Transcripts Alignment to a Reference) RNA-seq aligner v2.7.10b. Reads that were uniquely aligned to annotated genes were counted with featureCounts module of subread v2.0.5 [46]. Differential expression analyses were performed using DESeq2 v1.40.1 [47] using normal LFC shrinkage estimator for visualization and ranking purposes. Quality checks were performed on aligned BAM files and read count files using MultiQC v1.10.1. which gives a summary for all quality assessments.

### Gene set Enrichment and Gene Ontology analysis

GSEA v4.3.2 Mac App was downloaded from website (https://www.gsea-msigdb.org/gsea/) and annotated gene sets were used from Molecular Signature Database - MSigDB (v2022.1.Mm). We specifically focussed on biological, molecular and cellular pathways from m5.go.v2022.1, which is a list of ontology gene sets. GSEA analysis was performed using GSEAPreranked method, where genes were ranked by fold change and a ‘classic’ enrichment statistics was used to remove the magnitude bias of ranking metric. Minimum of 20 and maximum of 500 was defined as cutoff for number of genes in a gene set identification with maximum 1000 permutations. Network plots for GSEA categories were created using igraph_1.4.3 in R version 4.3.0 where the number of shared genes were represented as weights between the two points. GSEA comparison across datasets was performed for significant terms using a cutoff of Nominal *P* values (*P* <0.01) or FDR (*P*adj<0.1) as indicated and ranked by normalized enrichment score (NES). Tables summarizing these GSEA categories are supplied as supplementary datasets. Gene ontology analysis was performed using ClusterProfileR v4.12 [48] and enrichR [49]. For each analysis only significant terms were selected with a maximum *P* value <0.01 or as indicated.

### Transcript Length analysis

CA1 excitatory neuron specific dataset was retrieved from a publicly available data at GEO GSE74985. Transcript abundance was calculated using RSEM v1.3.0, where rsem-prepare-reference was used to extract reference transcripts and then rsem-calculate-expression to calculate the expression values. The transcript length and other genomic features were obtained from BioMart and the length of most abundant transcript was used for comparison of CDS length at gene level. Then the transcripts from translating mRNA (TRAP) and total RNA fraction of *Syngap^+/-^*, *FMR1^−/y^*, LTP and LTD datasets were separated in the bins of >1kb, 1-2kb, 2-4kb and>4kb and analysed for their up- and down-regulation.

### Percent Overlap estimation

For the percent calculation shown under the venn diagram overlaps, the percentage was calculated for *first mentioned dataset in first venn circl*e. The overlap number from this dataset was used as numerator while the selected population including overlap - from total (as indicated with *P* value or *P* adjusted value) was used as denominator and outcome was multiplied to 100. Thereafter, the percent for similarly and opposite regulation from overlap results was calculated taking the identified set as numerator and overlap as total – denominator.

### Statistical analysis

All statistics were performed using R. For RNA-seq datasets, differential expression was determined using DESeq2 using the default cutoff for significance (adjusted *P* value < 0.1). For GO and GSEA, significance was determined by nominal *P* value, and adjusted p-values with FDR cutoff where indicated. Differences between distributions were compared using two-sample *z* test as indicated.

## Data and materials availability

All transcriptomics sequencing raw data will be deposited in the Gene Expression Omnibus (GEO) upon publication. For analyses of cLTP, mGluR-LTD and *Fmr1^-/y^* TRAP-seq the following published datasets were used: GSE74985, GSE79790, GSE201239 and GSE101823.

## Author Contributions

EKO, SS, AS, and MR conceived the study and designed experiments. SS performed TRAP-seq isolation and initial bioinformatics analyses. AS and MR performed the majority of bioinformatic analyses. EKO wrote the manuscript with help from AS and MR.

## Acknowledgements and funding sources

The authors are supported by grants from the Wellcome Trust (219556/Z/19/Z), Medical Research Council (MR/S026312/1), H2020 Marie Skłodowska-Curie Actions (Syn2Psy) and Simons Initiative for the Developing Brain.

---

**Fig. S1.**
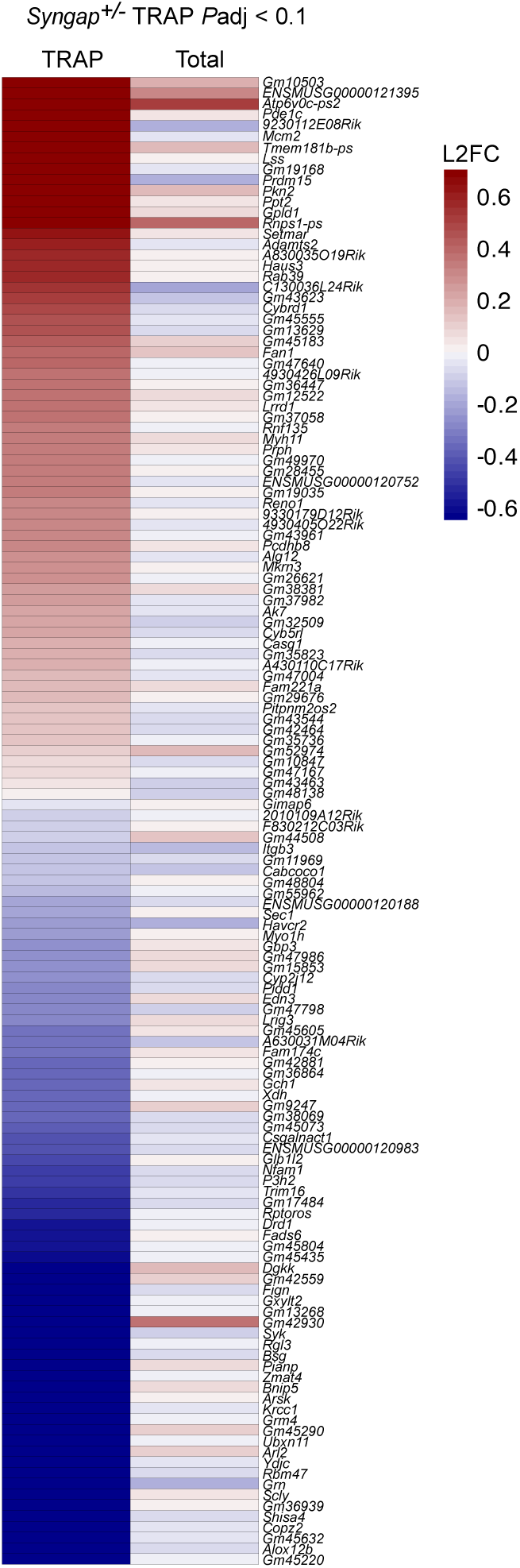
Significantly altered transcripts in *Syngap^+/-^*CA1-TRAP are not changed in total transcriptome. A heatmap of log2foldchanges (L2FC) shows that significant transcripts altered in *Syngap^+/-^* translatome are not changed similarly in the total transcriptome.

**Fig. S2.**
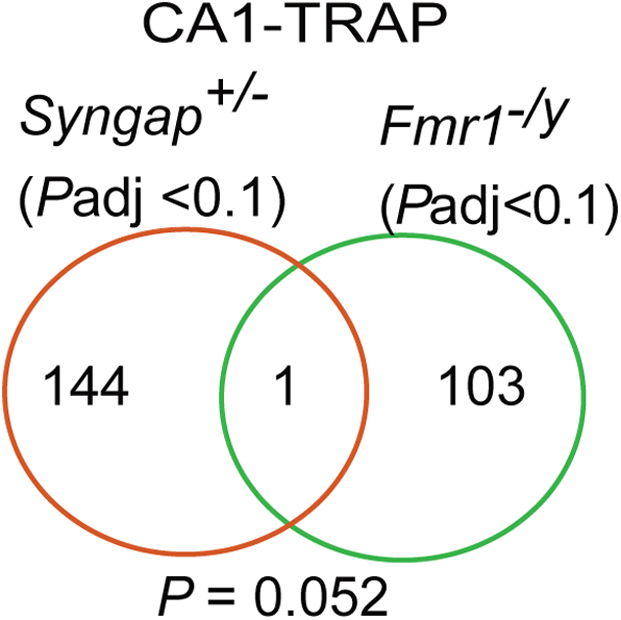
Mistranslation of distinct transcripts in *Syngap^+/^* and *Fmr1^-/y^*. Quantification of transcripts shows 145 significant transcripts (*P*adj < 0.1) are differentially translating in *Syngap^+/-^* and only 1 of those overlap with the significant translatome in *Fmr1^-/y^* (\**P* = 0.052).

**Fig. S3.**
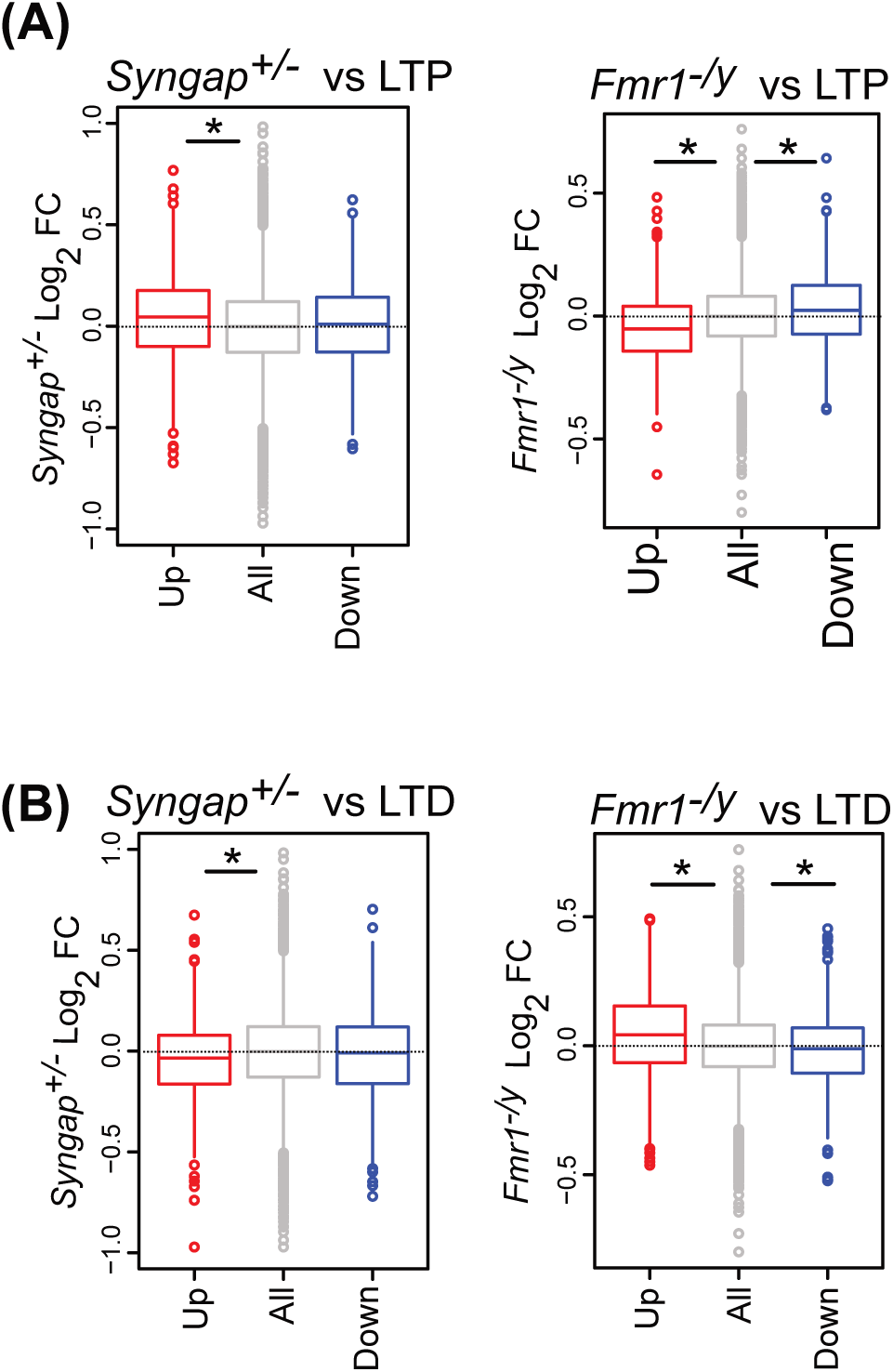
Comaprison of cLTP and mGLUR-LTD related translation changes in WT with *Syngap^+/-^* and *Fmr1^-/y^* CA1 neurons. **(A)** Analysis of the LTP significant transcripts in the *Syngap^+/-^*translatome shows significant increase in LTP- upregulated transcripts but no change in LTP- downregulated transcript (Kruskal-Wallis test \**P* = 3.30e-13, Post hoc two-sided Wilcoxon rank-sum test up \**P* = 4.77e-14, down *P* = 0.97), while LTP significant transcripts in the *Fmr1^-/y^* translatome show significant opposing change in both groups-LTP- upregulated and LTP- downregulated transcripts (Kruskal-Wallis test \**P* < 2.2e-16, Post hoc two-sided Wilcoxon rank-sum test up \**P* < 2.2e-16, down \**P* = 0.00013). **(B)** Analysis of the LTD significant transcripts in the *Syngap^+/-^* translatome shows significant decrease in LTD- upregulated transcripts (Kruskal-Wallis test \**P* = 9.81e-05, Post hoc two-sided Wilcoxon rank-sum test up \**P* = 2.29e-05, down *P* = 0.67). In contrast, LTD significant transcripts in the *Fmr1^-/y^* translatome show notably significant increase in LTD-upregulated transcripts (Kruskal-Wallis test \**P* =1.587e-11, Post hoc two-sided Wilcoxon rank-sum test up \**P* =7.71e-11, down *P* = 0.0026).

**Fig. S4.**
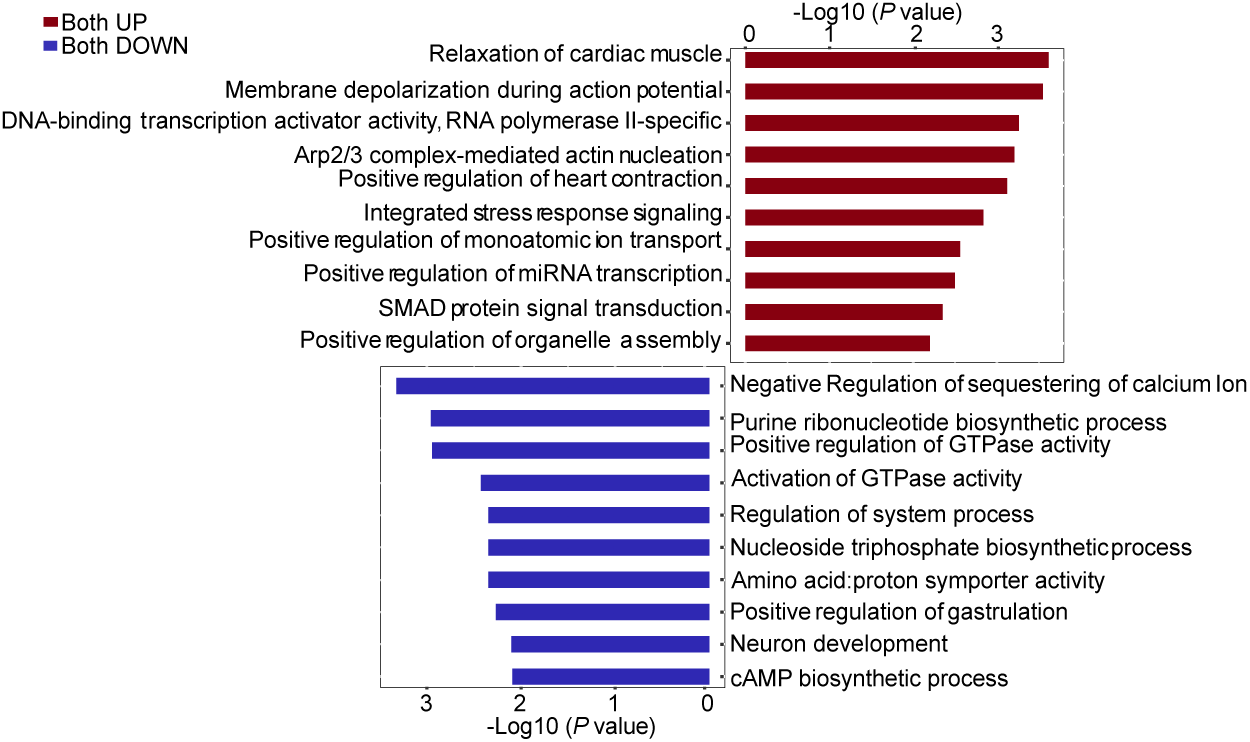
Gene ontology analysis of commonly changed transcripts in LTP and LTD. GO terms enriched in the overtranslated populations in both LTP & LTD regulate membrane depolarization, DNA binding while the commonly undertranslated transcripts primarily regulate calcium ion sequestration, GTPase activity, nucleotide biosynthesis.

**Fig. S5.**
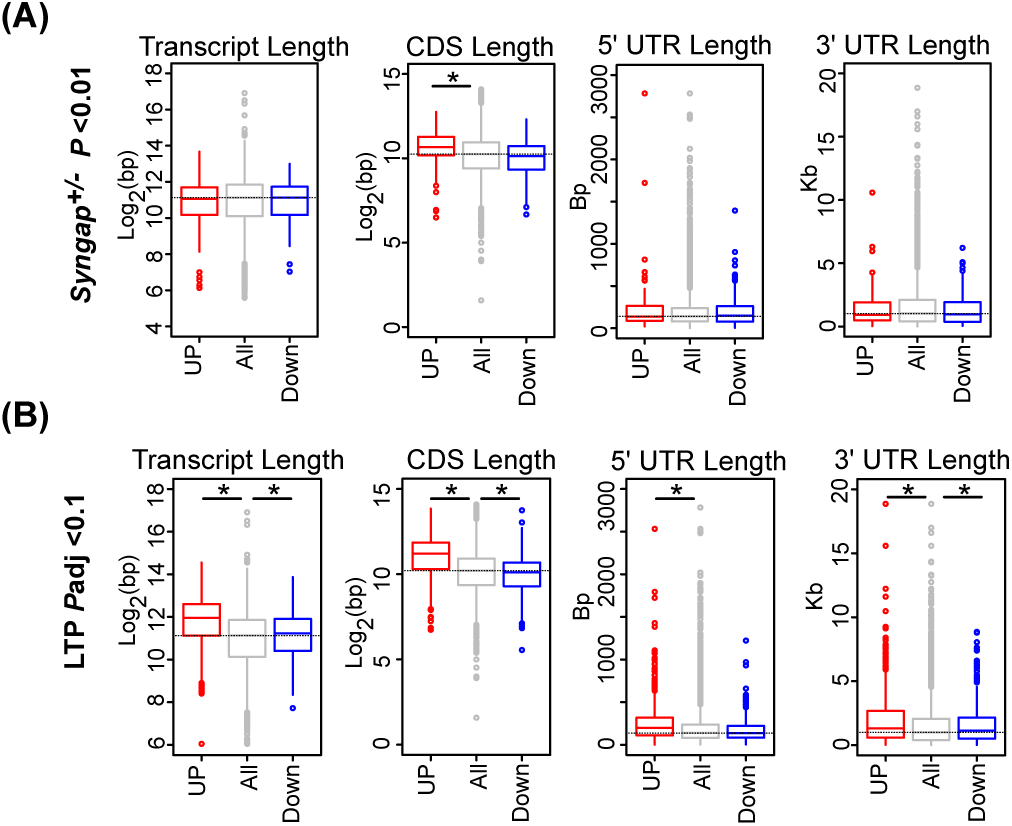
Genomic features analysis of significant transcripts (P value < 0.01) in *Syngap^+/-^* and cLTP datasets. **(A)** Analysis of the transcript length in *Syngap^+/-^* translatome (Kruskal-Wallis test *P* = 0.52); Analysis of the CDS length in *Syngap^+/-^*translatome (Kruskal-Wallis test \**P* = 0.0001, Post hoc two-sided Wilcoxon rank-sum test up \**P* = 3.80e-05, down *P* = 0.05); Analysis of the 5’ UTR length in *Syngap^+/-^* translatome (Kruskal-Wallis test *P* = 0.6362); Analysis of the 3’ UTR length in *Syngap^+/-^* translatome (Kruskal-Wallis test *P* = 0.88). **(B)** Analysis of the transcript length in LTP translatome (Kruskal-Wallis test *P < 2.2e-16, Post hoc two-sided Wilcoxon rank-sum test up \**P* < 2.2e-16, down \**P* = 0.017); Analysis of the CDS length in LTP translatome (Kruskal-Wallis test \**P* < 2.2e-16, Post hoc two-sided Wilcoxon rank-sum test up \**P* < 2.2e- 16, down \**P* = 0.002); Analysis of the 5’ UTR length in LTP translatome (Kruskal-Wallis test \**P* < 2.2e-16, Post hoc two-sided Wilcoxon rank-sum test up \**P* < 2.2e-16, down *P* = 0.15); Analysis of the 3’ UTR length in LTP translatome (Kruskal-Wallis test \**P* = 3.38e-09, Post hoc two-sided Wilcoxon rank-sum test up \**P* = 5.97e-10, down \**P* = 0.04).

## Additional files

**Additional file 1. *Syngap^+/-^* CA1 TRAP-seq.** DESeq2 results for all transcripts in TRAP-seq from hippocampal CA1 in *Syngap^+/-^* and WT littermates.

**Additional file 2. *Syngap^+/-^* total hippocampal transcriptome.** DESeq2 results for all transcripts in total hippocampal fraction from *Syngap^+/-^*and WT littermates.

**Additional file 3. GO analysis *Syngap^+/-^* CA1 TRAP-seq.** Gene ontology analysis terms for *Syngap^+/-^*CA1-TRAP significantly Up- and Down-regulated transcripts (*P*adj < 0.1) highlights nucleotide biosynthesis, protein modification and changes related to cellular resilience.

**Additional file 4**. **GSEA of *Syngap^+/-^* CA1 TRAP-seq.** GSEA on CA1-TRAP transcript population from *Syngap^+/-^*versus WT shows enriched gene sets related to DNA modification and recombinational repair.

**Additional file 5**. ***Syngap^+/-^*versus *Fmr1^-/y^*CA1-TRAP**. DEseq2 analysis of significantly differentially regulated transcripts in CA1-TRAP of *Syngap^+/-^*vs WT and *Fmr1^-/y^*vs WT.

**Additional file 6. GO analysis *Syngap^+/-^* versus *Fmr1^-/y^* CA1-TRAP.** Gene ontology analysis terms for *Syngap*^+/-^ CA1-TRAP significantly Up- and Down-regulated transcripts (*P* < 0.01) highlights DNA repair and shows minimal overlap with *Fmr1^-/y^*CA1-TRAP (GO terms.

**Additional file 7. GSEA of *Syngap^+/-^* versus *Fmr1^-/y^* CA1 TRAP-seq.** GSEA on the CA1-TRAP transcript population from *Syngap^+/-^* vs WT and *Fmr1^-/y^* vs WT reveals an overlapping population.

**Additional file 8. LTP-specific changes in Camk2a-ribotag population (*P*adj < 0.1)**. DEseq2 of Camk2a-ribotag transcripts from hippocampal slices at 30 min post-forskolin treatment (versus unstimulated).

**Additional file 9. GSEA comparison between *Syngap^+/-^*** /**WT, LTP/unstimulated, and *Fmr1^-/y^*/WT.** Comparison of gene sets changed in *Syngap^+/-^* TRAP, LTP ribotag, and *Fmr1^-/y^* TRAP.

**Additional file 10. LTD-specific changes in CA1-TRAP population (*P*adj < 0.1).** DEseq2 of CA1-TRAP transcripts from hippocampal slices at 30 min post-DHPG treatment (versus unstimulated).

**Additional file 11. GSEA comparison between *Syngap^+/-^*** /**WT, LTD/unstimulated, and *Fmr1^-/y^*/WT.** Comparison of gene sets changed in *Syngap^+/-^* TRAP, LTD TRAP, and *Fmr1^-/y^* TRAP.

**Additional file 12. Comparison between LTP- and LTD-specific changes (*P*adj < 0.1).** Commonly changed transcripts in both LTP and LTD datasets.

**Additional file 13. GO analyses of significantly upregulated transcripts in LTP and LTD datasets.** GO analyses of significantly upregulated transcripts in LTP-specific and LTD-specific datasets.

**Additional file 14. GSEA comparison between LTP/unstimulated and LTD/unstimulated datasets**. Comparison of GSEA enriched terms from LTP and LTD datasets.

**Additional file 15. Genomic features for *Syngap^+/-^*, *Fmr1^-/y^*, LTP, and LTD datasets.** List of Transcript length, CDS length, 3’UTR length and 5’UTR length of differentially expressed transcripts from *Syngap^+/-^*, *Fmr1^-/y^*, LTP and LTD datasets.

**Additional file 16. GO analyses of >2kb CDS transcripts significantly changed (*P* < 0.01) in LTP and LTD datasets**. Significant GO terms enriched in significantly changed >2Kb CDS transcripts from LTP and LTD datasets.

**Additional file 17. Convergent >2kb CDS transcript changes in *Syngap^+/-^* and LTP, and in *Fmr1^-/y^* and LTD populations**. List of >2kb CDS transcripts significantly upregulated in both *Syngap^+/-^*and LTP populations, and significantly downregulated in both *Fmr1^-/y^* and LTD populations.

